# A universal insect trait tool (ITT, v1.0) for statistical analysis and evaluation of biodiversity research data

**DOI:** 10.1101/2022.01.25.477751

**Authors:** Thomas Hörren, Martin Sorg, Caspar A. Hallmann, Vera M. A. Zizka, Axel Ssymank, Niklas W. Noll, Livia Schäffler, Christoph Scherber

## Abstract

We present a unique data set of trait information for 586 insect families in Central Europe, covering the largest known part of described species (over 34,000 species). Life history information and major functional traits were evaluated with fuzzy coding and weighted according to the number of known species in Germany. An overall analysis of the German insect fauna is given and the data set is exemplarily applied to metabarcoding results of malaise trap samples. Due to the high functional and taxonomic diversity in insects, further developments and refinements of traits to be included will be an ongoing process with advancements of upcoming database versions to be subsequently published.

## Introduction

Biodiversity research in terrestrial ecosystems needs a holistic perspective and a network approach to shed light on interspecific interactions determining the distribution and abundance of species and, ultimately, to understand causes and consequences of biodiversity change. Insects represent the most species-rich animal class in terrestrial ecosystems and have a tremendous effect on allmost all ecosystem processes. However, an overarching database compiling information about traits of most species or families is presently lacking. To understand interspecific interactions such as feeding relationships, it is necessary to take large data sets into account that encompass all families and species described so far for a particular area, rather than isolated information focusing on individual higher taxa. Such up-scaled trait data sets are paramount for the interpretation of all major monitoring efforts that rely on highly diverse composite samples, e.g. as those collected by malaise traps (Fig. 1 and 2). Lists of molecular units (OTUs - Operational Taxonomic Units or ESVs - Exact Sequence Variants) covering thousands of taxa retrieved by DNA metabarcoding without trait annotation, hinder any meaningful ecological interpretation of results. Previously, the only available approach to arrive at trait predictions for such data sets has been semantic language processing, a computation-intensive process potentially prone to misclassification. Here, we provide a trait-based assessment of Central European insect biodiversity covering the majority of currently described German insect species, which allows for retrospective and prospective assessments of changes in functional community composition.

**Fig. 1.**
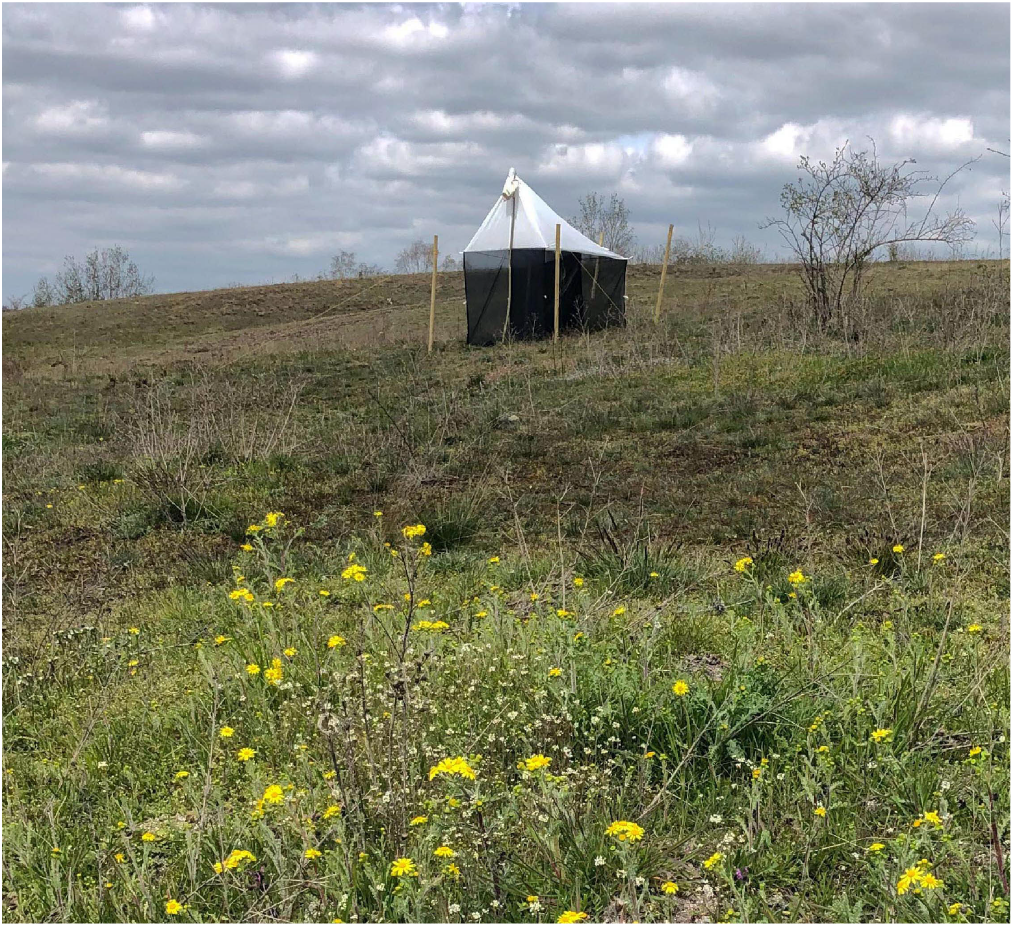
Malaise trap of the Townes model from the Entomological Society Krefeld in a German nature reserve.

**Fig. 2.**
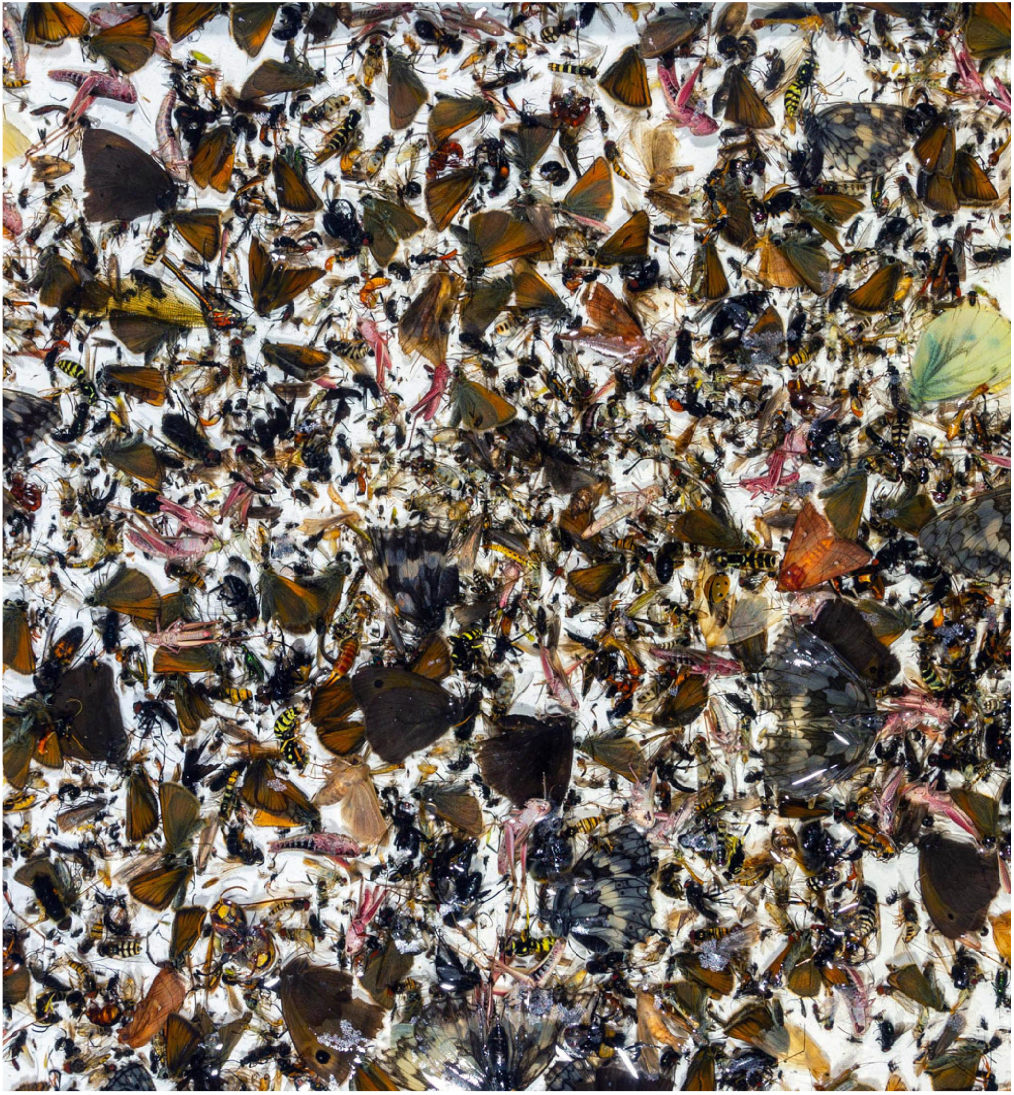
Bulk sample collected by a malaise trap of the Entomological Society Krefeld in a German nature reserve.

### Database development and components

Functional traits and behavioral preferences of species are increasingly used in ecology to understand interactions between organisms (e.g. https://opentraits.org/datasets.html). Our new insects trait tool (ITT) provides opportunity to analyze insect communities at family level and to assign trait values via proportions of absolute species numbers within a family. The ITT approach therefore compensates for incomplete connections between genetic barcodes and scientific species names and the presence of undescribed taxa. This methodical approach thus allows for handling complex insects taxa lists as those resulting from molecular biodiversity assessments and helps to understand and analyze the composition of insect communities.

The ITT contains trait information on 34,085 species from all of the 586 insect families occurring in Germany. The traits included so far focus on autecological feeding preferences of the larval stages. We focus on larval feeding ecology, as the larval stage reliably characterizes the habitat used for reproduction, and is better known than the diet of adult stages. The databases of the reference library “German Barcode of Life” (GBOL: https://bolgermany.de) served as reference base for total species numbers, and the Global Biodiversity Information Facility (GBIF: https://www.gbif.org) for individual queries in the process of developing the ITT. The systematic classification of the class Insecta in the GBOL reference library follows the definitions of Wheeler (1). We strictly follow this database here in order to take consistent reference data as a basis, even if the state of knowledge would entail revisions of the contents in some cases. The chosen approach in the ITT follows the method of ‘fuzzy coding’, as the expert assessment of biological information for the family taxa comes from different sources (2). We have compiled information from literature as well as field observations. The comprehensive literature sources are documented in Supplement 3. The classifications of zoophagous feeding types, in particular the classification of predation, micropredation, parasitism and parasitoid lifestyles follow the definition of Lafferty and Kuris (3).

All classifications made at family level are based on species- and genus-level traits that were categorized in 10% intervals within a system of decimal numbers ranging from 0 to 1.0. Smaller numerical proportions, e.g. terrestrial caddisfly larvae within the predominate larval aquatic insect order Trichoptera are not included in this resolution. Degrees of specialization, e.g. mono-, oligo- or polyphagy, have not yet been annotated, and there are uncertainties in categorization, i.e. unique feeding specializations such as in consumers of algae, moss or lichens that make a useful classification for comparative analyses difficult. Psocoptera, for instance, probably have highly specific feeding types, but as detailed information is lacking, they could only be classified as detritus feeders in the current version. Another example of higher taxa that still require classification by experts are Hemiptera (Auchenorrhyncha, Sternorrhyncha) and Thysanoptera, which have therefore also been grouped on a rather general basis.

Development of our insect trait database is to be considered a dynamic and ongoing process. Corrections and adjustments of ITT v1.0 will be continuously included in subsequent versions; in case of extensions or significant additions, a fully updated version of the database will be made publicly available.

### Table description, version 1.0

The matrix contains 29 columns in total. It lists all higher taxa at the level of order and suborder (column a) with the associated family taxon (column b) as well as the total number of species occurring in Germany and included in the “German Barcode of Life” (GBOL: https://bolgermany.de) (column c) in 586 lines of information. Columns t1-t26 contain the specified traits. All columns with traits contain decimal numbers from 0,1-1,0 within one group of columns counting 100% (each t1-t2, t3-t4, t5-t10, t11-t20, and t21-t26) which can be used as factors for each trait classification. Columns t1-t4 contain information on the link to aquatic (t1-t2) or terrestrial (t3-t4) ecosystems in the larval and adult stages. Columns t5-t10 contain information on the larval diet in numerical categories. These categories cover different organismic groups and decomposing material. Subsequently, there is a specification of the categories “phytophagous” and “zoophagous”. Phytophagous taxonomic families are broken down into feeding categories in columns t11-t20, zoophagous families in columns t21-t26. Supplement 1 contains the trait tables of the database in various file formats and a detailed explanation of the categories.

### Recommendations for use and examples

The insect trait tool refers to the insect fauna of Germany and is therefore predominately useful for taxa lists from Germany or the Central European region. This corresponds to an approximate area of application between the 47th and 57th degrees of latitude. Furthermore, the applicability is designed for comprehensive biodiversity samples as shown in Fig. 2. The ITT enables researchers to work with extensive taxa lists, even if those are not taxonomically standardized. The method is especially suitable for taxa-rich samples which are generated by certain examination techniques, for instance with malaise traps, car nets (4) or flushing samples from river banks (5). It is particularly useful for biodiversity assessments by metabarcoding when not all molecular units have yet been resolved to lower taxonomic levels, i.e. scientific names of genera or species remain unknown. This also concerns morphospecies in the case of classification by microscopic determination or artificial intelligence. The ITT table allows for a factor based classification of arthropod communities into aquatic and terrestrial habitats. In the central application area, larval feeding types can be assigned in the same way. Traits specified under “phytophagous” (t11-t20) and “zoophagous” (t21-t26) represent subordinate categories that should not be used for an overall consideration of communities contained in samples. However, these subordinate categories serve to further specify feeding behavior as if-then functions for taxa or groups of organisms with a primarily phytophagous or zoophagous lifestyle. The decimal numbers of individual traits can simply be used as factors for the resolved family, either directly to weight results or to relate them to the total species numbers in Germany. The latter is recommended when considering metabarcoding datasets and the number of taxa occurring in Germany still exceeds the number of taxa for which a barcode is deposited in reference libraries. Thus, with e.g. 67% coverage of barcodes of the real known number of species of a family in Germany, the result may be more realistic if this information is added as an correction factor.

We recommend the following steps for each application of the ITT:

1. Check your existing data set for suitability to apply the ITT. Your data set must have a reasonable size and should have its origin from the recommended region.
2. Harmonize the family systematics of your data set and the ITT. It does not matter which systematic is adjusted, it must be uniform.
3. An interpretation of your results can only take place within the information pattern of the ITT, as an approximation to reality.

### The insect diversity of Germany characterized with the trait tool

The last general overview of the insect fauna of Germany was published in 2003 (6) and further analyses have been based on the total number of 33,466 different insect species known back then. A comprehensive overview of ecological properties of all described species has not been published to date. The total lists generated from the GBOL databases during the development of ITT v1.0 cover 34,085 German insect species known to science. These species numbers within families were compared with the existing traits of the ITT. Due to the fact that the information from the specific feeding traits of the phytophagous (t11-t20) and zoophagous (t21-t26) groups are subsets of the basic diet types (t5, t6) there are deviations in the real species numbers. Calculated species numbers are a results of multiplication with often odd decimal numbers. They are therefore to be regarded as an interpretation. The relative distribution of species numbers is given in Fig. 3. Some patterns within an overall view are now recognisable for the first time. E.g. the proportion of aquatic species is 1.2 % for imagines and 8.7 % for larvae. This reflects the larval fixedness to aquatic habitats and allows the interpretation of high adaptation to mobility of imagines of most species. The most dominant major feeding types are zoophagous species with about 46,7% of the total species number and phytophagous species with about 36%. The further breakdown shows the specialisation of dependencies in the most species-rich organism group.

**Fig. 3.**
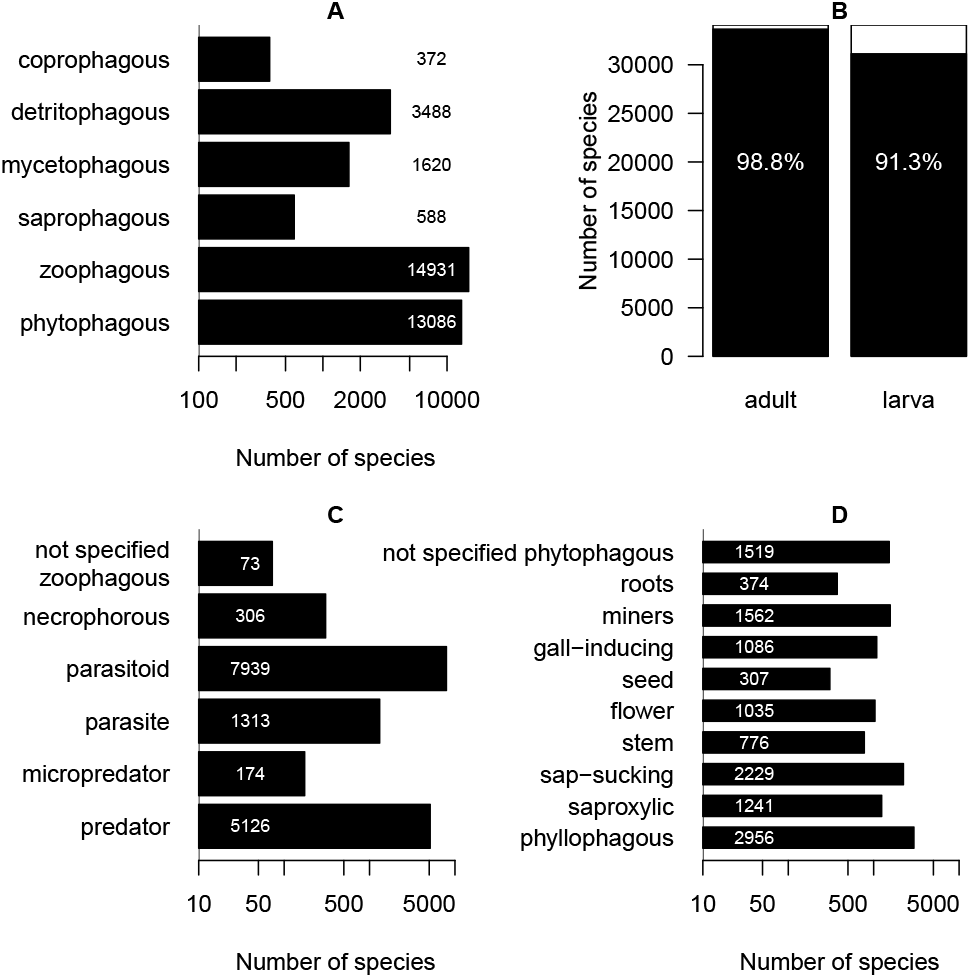
Distribution of trait categories for all known insect species in Germany. **A** Predominant diet types larvae. **B** Number of species associated with terrestrial habitat in larval and adult life stages. **C** Number of species in sub-classes among zoophagous species. **D** Number of species in sub-classes among phytophagous species.

### Evaluation of results from malaise trap samples

A standardized malaise trap of the Townes model from the Entomological Society Krefeld (7, 8) was used to measure insect diversity in the German nature reserve ‘Latumer Bruch’ near Krefeld at 51.326701N, 6.632973E. The study was carried out over the entire vegetation period in 2019 (March-October) with sample collection intervals of about 7-14 days. We selected five interval samples from late spring and summer (12-18 May, 29 May-8 June, 28 June-7 July, 7-18 July, 18-28 July) to analyze species rich seasonal insect communities with the ITT. Sequences yielded by metabarcoding were clustered into ‘Operational Taxonomic Units’ (OTUs) and taxonomic names were assigned by matching OTUs with reference databases (BOLD, GBOL and GenBank). The analysis (8) yielded 1,529 OTUs of arthropods, of which 1,355 were assigned to 163 insect families. The OTUs were summarized at the family level and the corresponding OTU table was combined with the ITT. Due to the inconsistent systematics in the family assignment, a harmonization of families was performed for the OTUs of each sample to calculate the share of larval and adult life form and larval feeding behavior in the examined community. The ITT was successfully applied to this metabarcoding results of highly diverse malaise trap samples. For the first time, the entire set of taxa contained in the samples could be analyzed at family level. At the same time, we conducted the most comprehensive analysis of trait information for malaise trap results. The distribution of the trait properties among the OTUs along considered samples can be found in Fig. 4. The results of the malaise trap samples vary within the trapping intervals. However, it is clearly visible that the patterns of the dominant groups are present even when looking at a short period and thus low species diversity (12-18 May) and similar to those of the longer intervals and thus higher species diversity (29 May-8 June, 28 June-7 July, 7-18 July, 18-28 July). Due to the late spring aspect and the summer aspect, no remarkable phenological patterns are recognizable. However, the May sample shows a higher overall proportion of gall-inducing insects. This fits with their high and species rich presence shortly after the beginning of the vegetation period.

**Fig. 4.**
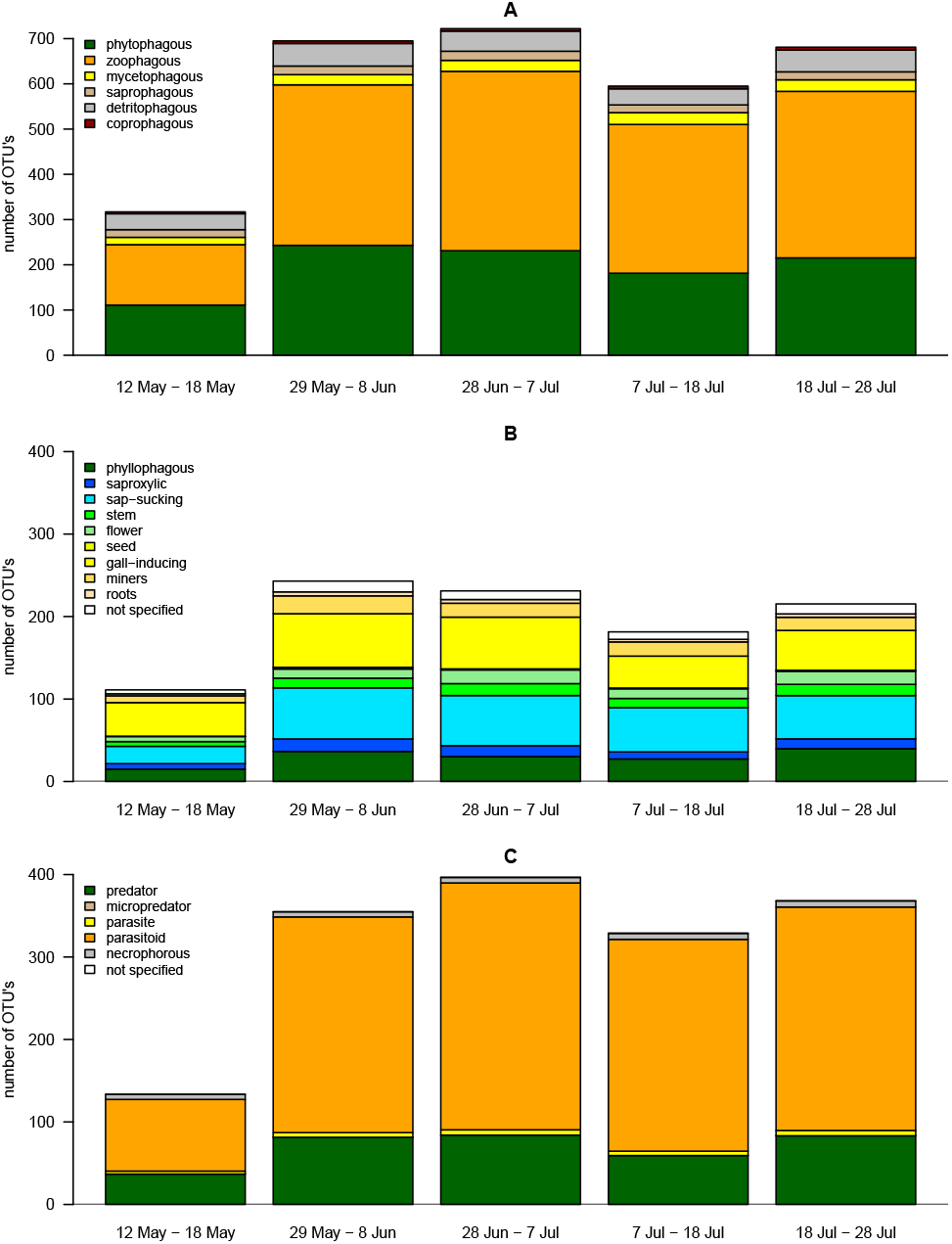
Distribution of major feeding strategies of insect OTUs along five consecutive malaise trap samples, using data from *Zizka et al*. (8). **A** Distribution of OTU numbers along major feeding strategies. **B** Distribution of OTU numbers in sub-classes among phytophagous species. **C** Distribution of OTU numbers in subclasses among zoophagous species.

In the species composition are the zoophagous (A) and parasitoid (C) taxa the dominant components along all five samples. Predators and parasitoids represent by far the largest proportions demonstrated in patterns.

Within phytophagous feeding types, which represent the second largest components, sap-sucking and thus the Hemiptera form the largest part within the captured phytophagous species (B). Overall, it is noteworthy that the distribution of traits assigned via the ITT in species-rich malaise trap samples resemble the distribution of dominant groups in the overall fauna of Germany (Fig. 3).

## Discussion

The insect trait tool (ITT, v1.0) is the first data set available for an overall trait assessment of Central European insect diversity. It is equally well suited for the analysis of insect communities in aquatic and terrestrial ecosystems. The ITT can be helpful to overcome identification difficulties associated with insect biodiversity studies.

The increase in knowledge is particularly significant for malaise trapping studies. There have already been species trait assesments in research combined with the application of standardized malaise traps used by the Entomological Society Krefeld, but only for partial analyses of selected groups (e.g. Hymenoptera Aculeata (9–11), Syrphidae (12), Lepidoptera (13), Trichoptera (14)).

First general, overall observations resulted in 1987 from the idea of measuring biomass information before sorting the insects contained in samples (15), a standardized method that also allowed for a comparative analysis in 2017 (16). However, the major limitation is still poor taxonomic knowledge and capacities, which is at least partially solved by DNA-based methods, particularly by metabarcoding (17). The detection of dominant structures in examples applied (Fig. 3 and 4) is particularly an interesting result since insects of several of these major feeding types play so far only marginal roles in conservation issues. This points to a serious weakness of many approaches and studies so far, because these taxa make up very large proportions of insect diversity. It concerns both interpretations within examined samples and the overall view of insect diversity. The ITT can therefore serve as an important tool in the analysis of insect diversity data sets for descriptive and applied research, the description of ecosystem functions and services, and nature conservation.

## Conclusions

The insect trait tool enables standardized comparisons of highly diverse insect communities with a minimum taxonomic resolution to family level. It is possible to compare patterns of samples, sites, relations to phenology, and correlative patterns in multivariate data sets (e.g. explanatory variables). It enables a comprehensive ecological evaluation of metabarcoding results or other comprehensive data sets like results from e.g. morphospecies approaches, artificial intelligence, and machine learning. An important advantage is that the ITT application compensates for lacking taxonomic knowledge and resolution, i.e. to include undescribed “dark” taxa of the Central European fauna and taxa for which so far no barcodes are assigned. In summary, the ITT can serve as an important tool for descriptive and applied research, the description of ecosystem functions and services, and nature conservation. Due to existing, massive deficits in the state of knowledge of the real overall diversity in the reference area, extended versions of this tool are required and intended in the future.

## ACKNOWLEDGEMENTS

Conceptual framework and development of methodologies of the EVK (TH, MS, CH) was funded by the German Federal Ministry for the Environment, Nature Conservation and Nuclear Safety (BMU), handled by the The German Federal Agency for Nature Conservation (BfN), grant number FKZ 3516850400. The metabarcoding analyses (VZ) were funded by the Ministry for Environment, Agriculture, Conservation and Consumer Protection of the German State of North Rhine-Westphalia (MULNV) (No. III-1-620.08). We would also like to thank many experts from the Entomological Society Krefeld for advice and assistance during the processing of the expert assessments.

## Author contributions

TH, MS and CH developed the conceptual framework. TH, MS and AS generated the traits and produced the trait database. VZ developed the examples from metabarcoding results. NN analyzed reference databases. CS contributed to traits and definitions. TH drafted the first version of the manuscript. All authors contributed to the article, approved the version to be published and agreed to be accountable for all aspects of the work.

**Supplement 1**

Insect trait tool (ITT) v. 1.0 database .pdf file: http://entomologica.org/tools/insect-trait-tool-v1-0.pdf

Insect trait tool (ITT) v. 1.0 database .xlsx file: http://entomologica.org/tools/insect-trait-tool-v1-0.xlsx

Insect trait tool (ITT) v. 1.0 database .pdf file.

Explanations of the trait characters: http://entomologica.org/tools/S1-explanations.pdf

**Supplement 2**

Application example for the insect fauna of Germany database .pdf file: http://entomologica.org/tools/insect-trait-tool-v1-0-fauna-germany.pdf

database .xlsx file: http://entomologica.org/tools/insect-trait-tool-v1-0-fauna-germany.xlsx

**Supplement 3**

Additional references used to classify lower taxa in the family trait categories as expert assessments: http://entomologica.org/tools/references-ITT-v1-0.pdf

